# No evidence that natural selection has been less effective at removing deleterious mutations in Europeans than in West Africans

**DOI:** 10.1101/002865

**Authors:** Ron Do, Daniel Balick, Heng Li, Ivan Adzhubei, Shamil Sunyaev, David Reich

**Affiliations:** Broad Institute of Harvard and MIT, Cambridge, MA, USA, 02142; Department of Genetics, Harvard Medical School, Boston, MA 02115; Division of Genetics, Brigham & Women’s Hospital, Harvard Medical School, Boston, MA, USA 02115; Howard Hughes Medical Institute, Harvard Medical School, Boston, MA, USA 02115

## Abstract

Non-African populations have experienced major bottlenecks in the time since their split from West Africans, which has led to the hypothesis that natural selection to remove weakly deleterious mutations may have been less effective in non-Africans. To directly test this hypothesis, we measure the per-genome accumulation of deleterious mutations across diverse humans. We fail to detect any significant differences, but find that archaic Denisovans accumulated non-synonymous mutations at a higher rate than modern humans, consistent with the longer separation time of modern and archaic humans. We also revisit the empirical patterns that have been interpreted as evidence for less effective removal of deleterious mutations in non-Africans than in West Africans, and show they are not driven by differences in selection after population separation, but by neutral evolution.

---

The effectiveness with which natural selection removes deleterious mutations from a population depends not only on the selection coefficient (*s*) of a mutation, but on the product of this and the population size (*N*) (*Ns*, reviewed in 1). Thus, the rate at which deleterious alleles are removed from a population depends on demographic history. Profound demographic differences across humans are well documented. Founder events in the last hundred thousand years have reduced the nucleotide diversity (the number of differences per base pair between an individual’s two chromosomes) in non-Africans by at least 20% relative to West Africans^2^^-^^4^, reflecting times when the ancestors of non-Africans had relatively smaller population sizes. Similarly, the advent of agriculture in the last ten thousand years has led to rapid population expansions.

To investigate the empirical evidence for natural selection having differed in its effectiveness across human populations, past studies have contrasted mutation classes thought to be subject to little selection (synonymous mutations in genes) to those potentially subject to purifying selection (non-synonymous mutations)^5^^-^^7^. The most important study on this topic measured the proportion of polymorphic positions in genes that are non-synonymous in 20 European and 15 African American samples, and showed that while all such segregating sites have a reduced rate in non-Africans, the reduction is proportionally less for non-synonymous sites.^5^ Based on simulations, this observation was interpreted as being consistent with reduced effectiveness of selection against weakly deleterious alleles in European than in West African populations due to their smaller size since separation^5^. Subsequent studies have confirmed the empirical observation and favored a similar interpretation^6^^,^^8^^,^^9^. These studies have clearly documented that there has been an interaction between the forces of natural selection and demographic history with regard to their effects on densities of non-synonymous sites^5^^,^^6^^,^^8^^,^^9^. However, such observations do not necessarily imply that there have been differences in the effectiveness of selection between the two populations after the split. An alternative explanation is that in the common ancestral population of Europeans and West Africans, the average derived frequency for non-synonymous alleles was lower than for synonymous alleles, as negative selection places downward pressure on the frequency of derived alleles^5^. The different initial distributions for the two classes of alleles would have responded differently to the bottleneck that then occurred in European populations, simply because they started out with different shapes, an effect that would be expected to occur even in the absence of any differences in the effectiveness of selection between the two populations since they split.

The most direct way to contrast the effectiveness of selection between two populations is to sample a single haploid genome from each population, count all the differences, and measure which of the two lineages carries an excess. Any two lineages that are compared in this manner must, by definition, be separated by the same amount of time since their common ancestor, and are thus expected to harbor the same number of lineage-specific mutations in the absence of selection or differences in mutation rate. In the presence of selection, however, mutations are removed from each lineage at a rate that depends on the product *Ns,* such that differences in the effectiveness of selection can be inferred from a detected asymmetry in the number of mutations in the two lineages. Here, we test for differences in the accumulation of mutations between two lineages by sampling one allele from population X and one from Y, determining the ancestral state based on the chimpanzee genome (*PanTro2*), and recording all the differences. We count the number of derived mutations in population *X* but not *Y* (*L*_*X not Y*_) and in *Y* but not *X*(*L*_*Y not X*_), and define a statistic *R*_*X/Y*_ = *L*_*X not Y*_ / *L*_*Y not X*_. We average over all possible pairs of samples, and compute a standard error using a Weighted Block Jackknife to correct for correlation among neighboring sites (Methods)^10^. If selection has been equally effective since the split and mutation rates have been the same on the two lineages, *R_X/Y_* should be within a few standard errors from 1.

We measured *R*_*WestAfrica/Europe*_ in four sequencing datasets: (1) coding regions of genes (exomes) from 15 African Americans and 20 European Americans^5^; (2) exomes from 1,089 individuals in the 1000 Genomes Project (1KG)^11^; (3) exomes from 1,088 African Americans and 1,351 European Americans^6^, and (4) 24 whole genomes sequenced to high coverage^12,13^ (Table S1). As expected for sites unaffected by selection and for mutation rates being indistinguishable on the two lineages, *R*_*WestAfrica/Europe*_ (*synonymous*) is always within 2 standard errors of 1 (Table 1 and Table S1). However, *R*_*WestAfrica/Europe*_(*non-synonymous*) is also indistinguishable from 1 (Table 1, Table 2, and Table S2). Thus, our empirical data provide no evidence that purging of weakly deleterious mutations has been less effective in Europeans than in West-Africans. This null result is consistent with the recently reported finding of ref.^14^ for similar population comparisons, which used a somewhat different approach. To generalize these results, we extended the analysis to a diverse set of human populations by computing *R*_*X/Y*_ between all possible pairs of 5 diverse sub-Saharan African and 6 non-African populations^13^, and fourteen 1000 Genomes Project populations. We observe no significant differences for any pair despite profound differences in demographic history (Table 2 and Table S3).

**Table 1.**
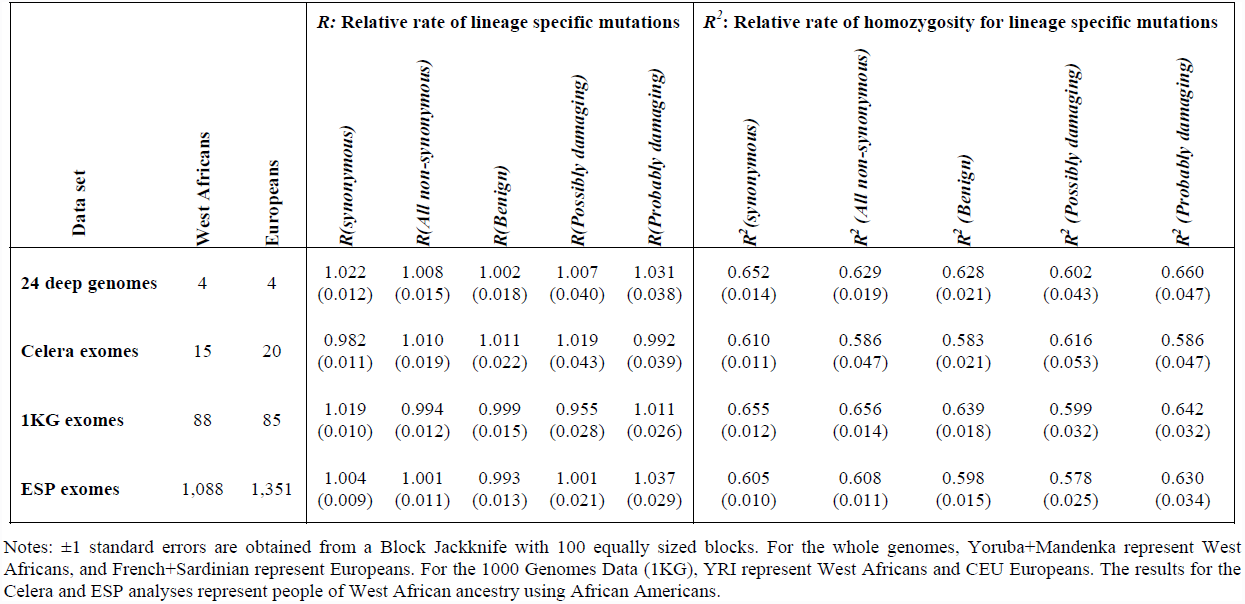
Accumulation of different classes of mutation in exomes of West African compared to exomes of European ancestry

| Data set | West Africans | Europeans | *R*: Relative rate of lineage specific mutations |  |  |  |  | *R*<sup>2</sup>: Relative rate of homozygosity for lineage specific mutations |  |  |  |  |
| --- | --- | --- | --- | --- | --- | --- | --- | --- | --- | --- | --- | --- |
|  |  |  | *R*(synonymous) | *R*(All non-synonymous) | *R*(Benign) | *R*(Possibly damaging) | *R*(Probably damaging) | *R*<sup>2</sup>(synonymous) | *R*<sup>2</sup>(All non-synonymous) | *R*<sup>2</sup>(Benign) | *R*<sup>2</sup>(Possibly damaging) | *R*<sup>2</sup>(Probably damaging) |
| **24 deep genomes** | 4 | 4 | 1.022<br>(0.012) | 1.008<br>(0.015) | 1.002<br>(0.018) | 1.007<br>(0.040) | 1.031<br>(0.038) | 0.652<br>(0.014) | 0.629<br>(0.019) | 0.628<br>(0.021) | 0.602<br>(0.043) | 0.660<br>(0.047) |
| **Celera exomes** | 15 | 20 | 0.982<br>(0.011) | 1.010<br>(0.019) | 1.011<br>(0.022) | 1.019<br>(0.043) | 0.992<br>(0.039) | 0.610<br>(0.011) | 0.586<br>(0.047) | 0.583<br>(0.021) | 0.616<br>(0.053) | 0.586<br>(0.047) |
| **1KG exomes** | 88 | 85 | 1.019<br>(0.010) | 0.994<br>(0.012) | 0.999<br>(0.015) | 0.955<br>(0.028) | 1.011<br>(0.026) | 0.655<br>(0.012) | 0.656<br>(0.014) | 0.639<br>(0.018) | 0.599<br>(0.032) | 0.642<br>(0.032) |
| **ESP exomes** | 1,088 | 1,351 | 1.004<br>(0.009) | 1.001<br>(0.011) | 0.993<br>(0.013) | 1.001<br>(0.021) | 1.037<br>(0.029) | 0.605<br>(0.010) | 0.608<br>(0.011) | 0.598<br>(0.015) | 0.578<br>(0.025) | 0.630<br>(0.034) |

Notes: $\pm 1$ standard errors are obtained from a Block Jackknife with 100 equally sized blocks. For the whole genomes, Yoruba+Mandenka represent West Africans, and French+Sardinian represent Europeans. For the 1000 Genomes Data (1KG), YRI represent West Africans and CEU Europeans. The results for the Celera and ESP analyses represent people of West African ancestry using African Americans.

**Table 2.**
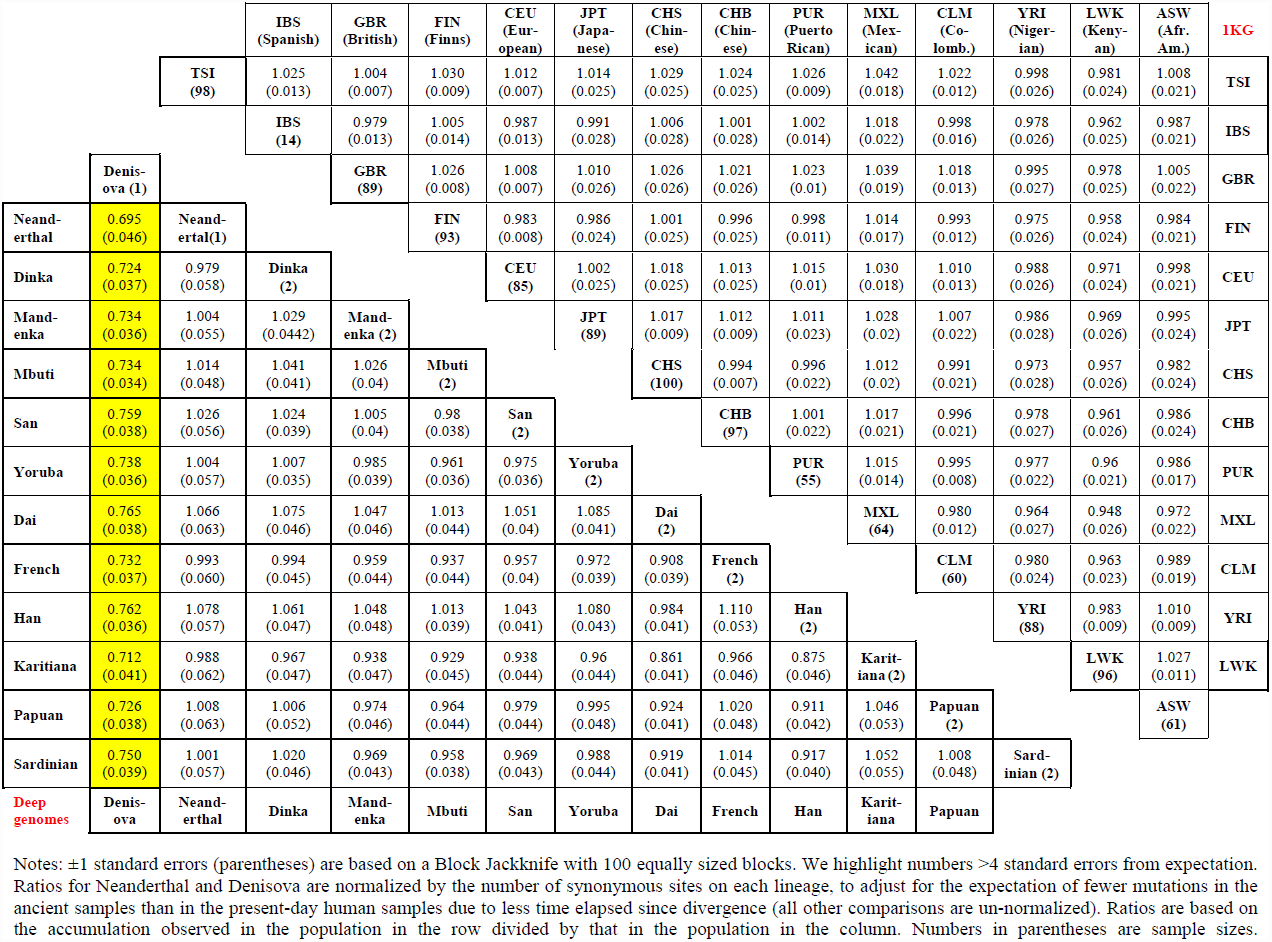
*R_X/Y_(probably damaging)* for all pairs of deep genome populations (bottom left) and 1000 Genomes populations (top right)

To interpret these null findings, we carried out simulations using fitted models of the demographic histories of West African and European populations^5^^,^^6^^,^^15^. (For most of our simulations, we assumed that mutations act additively although in Figure S1 we show that qualitatively different patterns are expected for mutations that act recessively, a phenomenon that we explore in a separate study^16^.) The simulations predict that *R_WestAfrica/Europe_* will be below 0.95 when *s* is between -0.004 and -0.0004 (Figure 1). However, if many mutations have selection coefficients outside this range, the signal could be diluted to the point of not being detectable. Indeed, when we compute the expected value of *R_WestAfrica/Europe_* integrating over a previously fitted distribution of selection coefficients^17^, we find that *R_WestAfrica/Europe_(non-synonymous)* is expected to be 0.987, too close to 1 to be easily detectable given the standard errors of our empirical measurements (Table 1)^18^. We next attempted to boost our power by stratifying non-synonymous sites based on their predicted functional effects. The PolyPhen-2^19^ and SIFT^20^ algorithms both predict function in a way that is dependent on the ancestral/derived status of allelic variants compared with the human reference genome, which has a particular ancestry at every segment that can bias measurements. We thus implemented a version of PolyPhen-2 that is independent of the allelic status of the human reference (Methods). We continue to observe no significant differences^18^ (Table 1, Table 2, Table S2, Table S3), although this null result may also reflect low power as when we infer the distribution of newly arising mutations for different PolyPhen-2 classes (Note S1), *R_WestAfrica/Europe_* is predicted to be 0.984-0.993 (Table S4).

**Figure 1.**
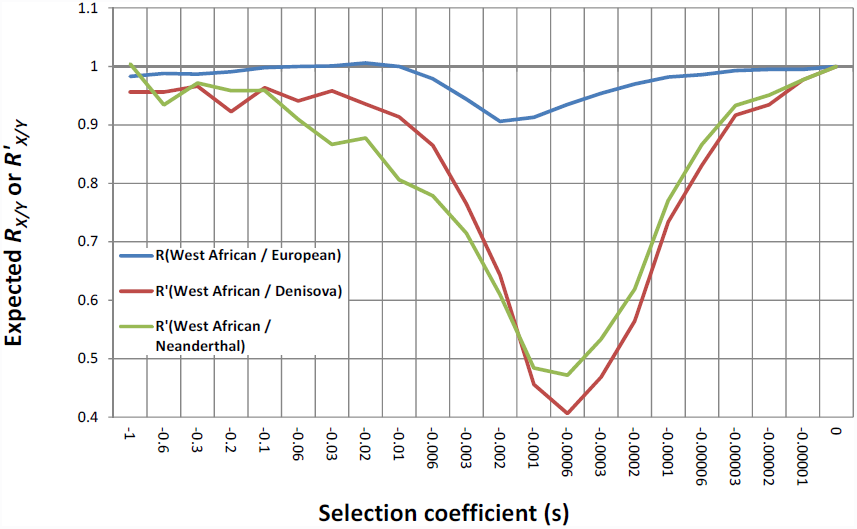
The effect of demographic history on the accumulation of deleterious mutations. To study the expected value of *R_Westafrica/Europe_* stratified by selection coefficient, we simulated a previously published model of the joint history of West Africans and Europeans^6^, for a range of selection coefficients, assuming an additive model of nature selection (Figure S1 shows similar results for other demographic models). The simulations show that *R_WestAfrica/Europe_* dips below a potentially detectable ratio of 0.95 for *s* ∈ (−0.0004, −0.004). We also simulated a published model of the history of archaic Denisovans, archaic Neanderthals, and West Africans ^13^. The simulations predicts similar curves for *R′_WesAfrica/Denisova_* and *R′_WestAfrica/Neanderthal_* reflecting their similar inferred demographic histories (we use a normalized *R′* statistic to correct for the effects of branch shortening in these ancient lineages). The simulations show that *R′_WestAfrica/Denisova_* is expected to below a detectable ratio of 0.95 for *s* ∈ (−0.00002, −0.03) and that *R′_WestAfrica/Neanderthal_* is expected to be below 0.95 for *s* ∈ (−0.00002, −0.09).

To increase statistical power to detect any real differences in the net effectiveness of selection, we leveraged the fact that the population split between African and non-African populations occurred only in the last roughly one hundred thousand years. Mutations that arose prior to the population divergence are expected to dilute any true signal. We therefore computed a time-stratified *R_African/non-African_*-statistic, taking advantage of data from 4 experimentally phased African and 6 experimentally phased non-African genomes, all processed similarly^13^. We created pseudo-diploid genomes by merging 1 African and 1 non-African phased genome, for a total of 96 = (4×2)×(6×2) comparisons, and then used the Pairwise Sequential Markovian Coalescent method (PSMC) to infer the local time since the most recent common ancestor (TMRCA) at each location, masking out the exome to avoid circularly using the same sites for inferring TMRCA and computing *R_African/non-African_*. Table S5 shows that when we restrict to the subset of the genome with the lowest inferred TMRCA in this analysis, we still detect no differences.

As a third way to increase statistical power, we analyzed a class of sites that is much larger than the class of coding substitutions so that we can make measurements with much smaller standard errors; this is the class of sites affected by biased gene conversion (BGC). BGC is a selection-like process in which DNA repair acting on heterozygous sites in gene conversion tracts favors GC over AT alleles^21^. Since BGC only acts on heterozygous sites, it occurs at a rate proportional to heterozygosity, or 2*p*(1-*p*) for a mutation of frequency *p*, exactly mimicking additive selection^21^. We find that 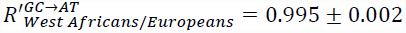 and 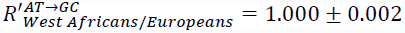 by using an 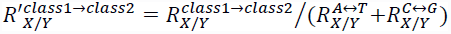 statistic that normalizes by the rate of A↔T and C↔G substitutions not expected to be affected by BGC. A further advantage of this normalization is that it also corrects for possible differences in mutation rates across lineages. For diverse population comparisons, we detect no differences significant at |Z|>3 standard errors from zero with the exception of San Bushmen who have about 1% more GC→AT mutations than other humans (significant at up to 8 standard errors) (Table S6). This is the first direct detection of less effective removal of mutations in some present-day humans than in others, and is consistent with the San being amongst the most deeply diverged human populations, which may have provided more opportunity for slight differences in the effectiveness of removal of mutations across populations to have a cumulatively measurable effect^22^.

To demonstrate empirically that differences in the accumulation of non-synonymous sites can be empirically detected given a sufficiently ancient population divergence time and sufficiently different subsequent demographic histories, we also analyzed two deeply sequenced genomes from archaic humans: Denisovan and Neanderthal. The ancestors of both are inferred to have maintained relatively small effective population sizes for on the order of a half million years since their main separation from present-day humans, consistent their levels of genetic diversity being 3-6 times smaller^12^. A challenge in comparing the accumulation of mutations in present-day human samples to ancient samples is that fewer mutations are expected to have occurred on the lineage of ancient samples because they are from individuals who lived closer in time to the common ancestor. To correct for this, we divide the accumulation of non-synonymous mutations on each lineage by the synonymous sites: *R′_X/Y_(non-synonymous class)* = *R_X/Y_ (non-synonymous class)*/*R_X/Y_(synonymous)*. After removing C→T and G→A mutations in the archaic genomes that have evidence of degradation leading to ancient DNA errors, we find that present-day humans have accumulated deleterious mutations at a significantly lower rate than Denisovans since separation: *R′_Modern/Denisova_(non-synonymous)* = 0.872 ± 0.034 (P=0.0002) (Table S7)^12^. In contrast, *R′_Modern/Neanderthal_(non-synonymous)* = 1.037 ± 0.037 is consistent with 1, suggesting that deleterious mutations have not been removed as effectively on the Neanderthal as on the Denisovan lineage (Table S7). The higher resolution BGC analysis further detects that the effectiveness of removal of mutations on the Neanderthal lineage was intermediate between that of Denisovans and modern humans: 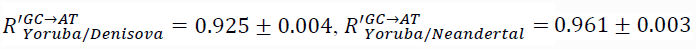, and 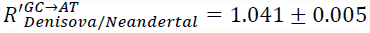. The different rates of accumulation in Neanderthals and Denisovans despite similar inferred demographic histories (Figure 1) suggests that fitted models of demographic history, e.g. for West Africans and Europeans, may not always provide accurate predictions of the relative effectiveness of removal of mutations.

In light of these results, is there any reason to believe that weakly deleterious mutations have been removed less effectively in Europeans than in West Africans? The strongest previous evidence for such an effect was based on an alternative statistic: the proportion of polymorphic sites in the exome that are non-synonymous^5^. We therefore carried out simulations of demographic history that allowed us to study the change in this statistic over time; our simulations agree with previous simulations of the same statistic^5^, but allow for additional insights into the evolutionary forces responsible for the dynamics (Figure 2). In the first set of simulations, we adjusted the selection coefficient *s* in Europeans in each generation so that the quantity governing the effectiveness of selection *Ns* was by construction always the same in Europeans and West Africans. The simulations show that the proportion of non-synonymous sites in Europeans in fact rises more, not less, when we adjust our simulations in this way to eliminate any differences in the effectiveness of selection (blue curves in Figure 2B). Thus, the observed rise in the proportion of non-synonymous sites in Europeans is not due to differences in the effectiveness of selection after the population split, but occurs in spite of such differences. In the second set of simulations, we partitioned the change in the proportion of non-synonymous sites over time into selection and neutral effects. The simulations show that the dynamics of the previously studied statistic are driven by neutral forces (correlation coefficient of ρ = 0.96 to 0.99). In contrast, changes in the effectiveness with which selection removes deleterious alleles in Europeans compared with West Africans has an effect that is opposite to the observed change (ρ = -0.45 to -0.27) (Figure 2C and Figure 2D). Intuitively, prior to the West African / European split, allele frequencies of non-synonymous polymorphisms would, on average, have been much lower due to the depletion of non-synonymous sites by selection, and the per-site density of non-synonymous segregating sites would also have been lower. The population entering the bottleneck primarily loses rare alleles, so the non-synonymous site allele frequency distribution would be expected to adjust faster than that for synonymous sites. Once the population re-expands, the allele frequency distribution for non-synonymous sites also adjusts faster, in this case because the same flux of new mutations into both classes causes a faster rate of replenishment of non-synonymous sites than synonymous sites due to an initially lower density. These findings illustrate the complexity of the interactions between selection and demographic history in their effects on genetic variation, and highlight an opportunity first identified by ref. ^5^, which is that joint analyses of demographic history and natural selection can provide more insight into the nature of both evolutionary forces that either type of analysis alone.

**Figure 2.**
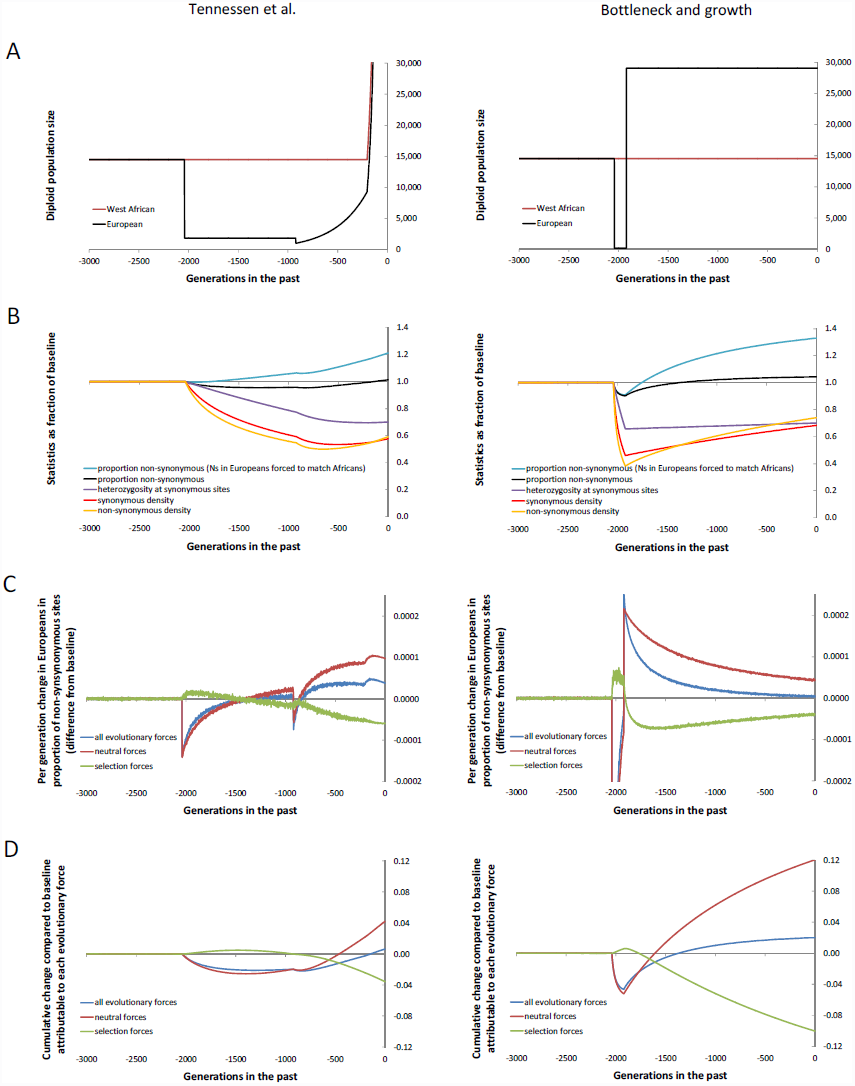
The rise in the proportion of non-synonymous sites in Europeans compared with West Africans is not due to a reduced effectiveness of selection in Europeans since the split. (A) The West African and European diploid population sizes for the two simulated models, both of which specify a population split 2,040 generations ago. The plots of the temporal dynamics of the statistics in subsequent panels are restricted to the European population, as the West African population size does not fluctuate enough to result in appreciable changes from the baseline. (B) We show the values of key statistics as a fraction of the baseline. The present-day proportion of non-synonymous sites in Europeans is higher than in the ancestral populations (black curves). This cannot be attributed to less effective removal of deleterious mutations in Europeans than in Africans since the population split, as we can see from the fact that when we carry out simulations in which the selection coefficient in Europeans per generation is set so that the effectiveness of selection *Ns* in the two populations is the same, the rise actually becomes greater (blue curves). (C) Partitioning of the change in the proportion of non-synonymous sites per generation into neutral and selective forces shows that for the West African / European comparison, the temporal dynamics are driven by neutral forces (strong positive correlation in the dynamics) and not by the selective forces (negative correlation). (D) Plots of the cumulative effect of each evolutionary force compared to baseline show that differences in the effectiveness of selection between Europeans and West Africans since they separated in fact have a net negative, not a net positive effect on the proportion of non-synonymous sites in Europeans.

It is tempting to interpret the indistinguishable accumulations of deleterious mutations across present-day humans as implying that the overall genetic burden of disease should be similar for diverse humans. To the extent that mutations act additively this is correct, and it implies that the complex demographic events of the past are not expected to lead to substantial population differences in prevalence rates of complex disease that have an additive genetic architecture^18^. However, recessively or epistatically acting mutations work in combination to contribute to disease risk, and since demographic history affects allele frequencies, it affects the rate of co-occurrence of alleles. For example, Table 1 and Table S8 show that the absolute count of alleles occurring in homozygous form is empirically higher in non-Africans than in Africans for all functional site classes, confirming previous findings^5^. Thus, the relative risk for diseases that are contributed to by recessively acting mutations could be expected to be influenced by demography, as indeed is known to be the case for populations that experienced recent founder events like Ashkenazi Jews and Finns. An important direction for future work is to determine the extent to which mutations contributing to phenotypes act non-additively, as this will determine the extent to which demographic history is important in affecting human disease risk.

## Methods

### Data

The datasets we analyzed were published previously and are summarized here.

“*Celera*”: PCR amplification and Sanger sequencing was performed on 15 African American (AA) and 20 European American (EA) samples over the coding sequences of 10,150 genes. We downloaded ancestral and derived allele counts for 39,440 autosomal SNPs from the supplementary materials of the original study, restricting to sites with genotypes available from both AA and EA^5^.

“*1000 Genomes (1KG)*”: A total of 1,089 samples from 14 populations were analyzed in Phase 1 of the 1000 Genomes project. Illumina-based exome sequencing^11^ was performed to ~100× average coverage after solution hybrid capture of the exome^23^.

“*ESP*”: A total of 1,088 African Americans and 1,351 European Americans were sequenced as part of the National Heart, Lung and Blood Institute Exome Sequencing Project. Illumina-based exome sequencing was performed to ~100× average coverage after solution hybrid capture of the exome ^6^.

“*24 Genomes*”: This dataset includes 2 samples each from 6 non-African and 5 sub-Saharan African genomes, an archaic human from Denisova Cave in Siberia sequenced to 30× coverage, and an archaic Neanderthal from Denisova Cave in Siberia sequenced to 52× coverage. All sequencing data is based on Illumina technology. We used the version of this dataset reported in ref. ^13^. We only analyzed sites with genotype quality scores (GQ) of ≥45^24^.

### Mutation annotation

We annotated coding mutations using ANNOVAR^25^, which classifies sites as “non-synonymous”, “synonymous”, “stop-gain” or “stop-loss”. We sub-classified variants using a simplified version of PolyPhen-2 that is independent of the ancestral/derived status of the human genome reference sequence (“human-free Polyphen-2”). To guarantee independence of PolyPhen-2 predictions from the human genome reference sequence, we created a simplified version that relies solely on the multi-species conservation score used in this method^26^. This score reflects the likelihood of observing a given amino acid at a site conditional on the observed pattern of amino acid changes in the phylogeny, and is the most informative feature of PolyPhen-2. The predictions in our simplified PolyPhen-2 method are based on the absolute values of the difference of the scores for the two alleles. By construction, this is symmetric with respect to reference/non-reference (and also ancestral/derived, major/minor) allele status. This procedure is similar to the original version of PolyPhen, but relies on the PolyPhen-2 homology search and alignment pipeline.

### Statistics

We are interested in the expected number of mutations in a randomly sampled haploid exome from one population that are not seen in a randomly sampled comparison exome from another population. To compute this in a situation where we have many exomes available from each population, we do not wish to literally randomly choose an exome from each population as this would throw away data decreasing the precision of our estimates. Instead, we obtain the expected value if we performed an infinite number of random samplings. To compute this, at each variable site *i* in the genome we define 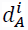 as the count of the mutant allele at that site in a sample of 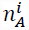 exomes from population A and 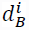 as the count of the mutant allele in a sample of 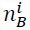 exomes from population B. The expectations are obtained by summing over all sites in the genome.

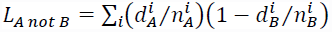

For some analyses, we also wished to compute the relative probability that a population is homozygous for a derived allele whereas the other is not. Thus, we defined two additional statistics, now imposing a correction for limited sample size (since we need to sample two alleles from each population, we need to sample without replacement):

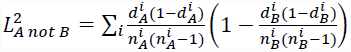

We then defined the ratio statistics

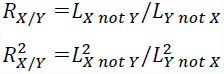

### Weighted block jackknife to estimate standard errors

We obtained standard errors using a weighted block jackknife^10^. We divided the SNP datasets into 100 contiguous blocks and then recomputed the statistic on all the data except for the data from that block. The variation can be converted to a standard error using jackknife theory. We assess significance based on the number of standard errors from the null expectation of R=1, and compute a P-value using a Z-score assuming a normal distribution.

### Time-stratified computation of relative accumulation of deleterious mutation

We began with data from 10 experimentally phased genomes, all processed in a nearly identical way^13^. These genomes consisted of one each from the populations in Table 2 except for the Dinka. We then combined haploid genomes from one of 4 African and one of 6 non-African individual to make 96 = (2×4)×(2×6) pseudo-diploid individuals. We masked the data from the exome, and ran the Pairwise Sequential Markovian Analysis (PSMC)^2^ on the data to estimate the time since the most recent common ancestor of the two phased genomes at each location in the genome. We stratified the data into three subsets of inferred time depth, and then computed the *R_African/non-African_*-statistic within each time-stratified subset (using exomic sites that had been masked from the PSMC analysis so we could independently use them for analysis).

### Analysis of sites susceptible to biased gene conversion (BGC)

We computed the accumulation of mutations susceptible to BGC for three different classes: GC→AT mimicking negative additive selection, AT→GC mimicking positive additive selection, and A/T or G/C polymorphisms which we treat as neutral (and use as the denominator of *R′_X/Y_*). For BGC analyses we use the entire genome, after excluding sites in the exome. For analyses involving ancient samples, we exclude C→T and G→A sites from the analysis of GC→AT substitutions (we restrict to C→A and G→T substitutions), to avoid the degradation errors typical of ancient DNA that we are concerned may affect even the consensus genome sequences.

### *R′_X/Y_*-statistic: Correcting for branch shortening and differences in mutation rates

For analyses involving the archaic Denisovan and Neanderthal samples, which are many tens of thousands of years old and thus have experienced less evolution from the common ancestor than present-day humans to which they are compared, we do not expect that *L_archaic not modern_ = L_modern non archaic_* even for neutral sites. For all analyses involving ancient samples, we instead compute normalized statistics *L_A not B_′* and *L_B not A_′*, where we divide both *L_A not B_* and *L_B not A_* by the accumulation of mutations at sites that are expected to act neutrally (synonymous sites for coding sequences and A/T + C/G for BGC). Thus, 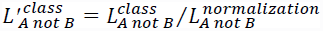. We then define:

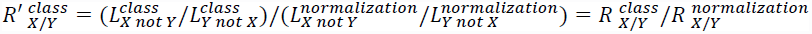

This *R′_X/Y_*-statistic not only corrects for branch shortening in the ancient samples, but also has the benefit of correcting for any differences in mutation rate that might have arisen on one lineage or another since population separation.

### Simulations

We wrote a forward simulation in C that implements an infinite sites model. Each mutation is assumed to occur at an unlinked site. To the extent that linkage affects the expected values of the statistics we compute, our simulations are not capturing these effects.

There is an initial burn-in period of 250,000 generations to generate an equilibrium allele frequency spectrum. The simulator samples the allele counts in the current generation based on the frequencies in the previous generation, the selection coefficient *s*, the dominance coefficient *h* (usually set to additive or *h*=0.5), and the current population size.

For modeling West African and European history in the simulations reported in the main text, we use a demographic model previously fitted to genetic data^6^. (Figure S1 reports results for other four different demographic histories, shown in Table S9). For comparisons of West African and archaic population history we also use a previously fitted demographic model^13^. We use a mutation rate of 2×10^−8^/base pair/generation.

For analyses of the proportion of sites that are expected to be non-synonymous in a sample size of 40 chromosomes, we use a hypergeometric distribution to obtain the expected value of this statistic. If the population size in a generation is *N* and *K_i_* is the number of copies of the derived allele at site *i*, then we can compute the probability that a sample of 40 chromosomes is polymorphic at a site as 1 minus the hypergeometric probability of 0 or 40 derived alleles:

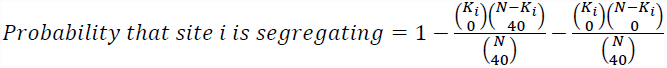

We average this probability over all simulated positions to obtain the density of segregating sites.

### Integrating over distributions of selection coefficients

For some statistics, we wished to obtain an expected value integrating over distributions of selection coefficients. To achieve this, we carried out simulation series for different selection coefficients; for example, for Figure 2, each of 19 values: *s* = { −1×10^6^, −2×10^6^, −5×10^6^, −1×10^5^, −2×10^5^, −5×10^5^, −1×10^4^, −2×10^4^, −5×10^4^, −1×10^3^, −2×10^3^, −5×10^3^, −0.01, −0.02, −0.05, −0.1, −0.2, −0.5, −1}. To compute expected values for *L_X not Y_*, *L_Y not X_*, and the density of segregating sites per base pair in a fixed sample size of 40 chromosomes, we use a weighted average of the values of the simulated single selection coefficient statistics. For some analyses, we use the distribution of human selection coefficients for non-synonymous sites from ref. ^17^, where the probability of a given value of -*s* is drawn from a gamma distribution fitted to European genetic data with α=0.206 and β=15400. For analyses of the expected value of *R_WestAfrica/Europe_* stratified by PolyPhen-2 functional class, we use the values inferred in Note S1.

### Simulations forcing the effectiveness of selection in Europeans and West Africans to be the same

To evaluate whether differences in the effectiveness of selection in the Europeans and West African populations since their split could be explaining the observed rise in the proportion of non-synonymous sites in Europeans above baseline, we modified the simulator so that in every generation *i*, the selection coefficient in Europeans *s_e,i_* is determined dynamically. Define *N_e,i_* and *N_a,i_* as the diploid population sizes in Europeans and Africans, respectively, and define the selection coefficient in Africans (held constant) as *s_a,i_.* We then set the selection coefficient in Europeans in each generation to be *s_e,i_* = *s_a,i_(N_i,a_/N_i,e_*). This procedure has the consequence that *N_e,i_S_e,i_ = N_e,i_S_a,i_(N_a,i_/N_e,i_) = N_a,i_S_a,i_*. Since the quantity *Ns* governs the effectiveness of selection, selection is guaranteed to be equally effective in both populations at all times.

### Partitioning the evolutionary dynamics into the effects of selection and neutral effects

We modified the simulation to sample two counts of derived alleles in each generation for a given selection coefficient *s* and nucleotide position *i*. The first count is *A_s,i_*, which reflects the forces of selection, mutation, and stochastic sampling. The second is *N_s,i_*, which reflects the effects of mutation and stochastic sampling but not selection. The counts in each generation are only updated based on *A_s,i-1_* in the previous generation, so the long-term evolutionary trajectory properly incorporates the effects of natural selection as they have accumulated over generations.

We average the values of *N_s,i_* and *A_s,i_* over simulation replicates, and compute a weighted average of the results based on the distribution of selection coefficients from ref.^17^. We define the proportion of sites that are expected to be non-synonymous in a given generation as *PropAll_i_* = *A_s,i_/(A_s,i_+ A_0,i_*), and the proportion that would be non-synonymous if selection had been switched off in that generation as *PropNeu_i_* = *N*_s,i_*/(N_s,i_+ A_0,i_*).

With this notation, the expected change in the proportion of nonsynonymous sites due to all evolutionary forces in generation *i* is *δPropAll_i_ = PropAll_i_- PropAll_i_*. The proportion due to neutral forces only is *δPropNeu_i_* = *PropNeu_i_- PropAll_i-1_,* and the proportion due to selective forces only is *δPropSeli* = *PropAll_i_- PropNeu_i_*.

To compare the effect of an evolutionary force in a given generation to what it was in the ancestral population prior to the split (>2,040 generations ago in the simulations of Figure 2), we defined the baseline-corrected statistics *ΔPropSel_i_* = *δPropSel_i_-δPropSel_baseline_, ΔPropNeu_i_* = *δPropNeu_i_-δPropNeu_baseline_*, and *ΔPropAll_i_* = *δPropAll_i_-δPropAll_baseline_*. These statistics are positive if the effectiveness of removal of mutations due to an evolutionary force is less than in the ancestral population, and negative if the effectiveness is greater than in the ancestral population. These are the statistics plotted in Figure 2C and integrated to obtain cumulative effects in Figure 2D.

## Acknowledgments

We are grateful to Joshua Akey, David Altshuler, Carlos Bustamante, Sergi Castellano, Cesare de Filippo, Alexey Kondrashov, Eric Lander, Kirk Lohmueller Swapan Mallick, Svante Pääbo, Nick Patterson, Jonathan Pritchard, Joshua Schaiber, Guy Sella and Montgomery Slatkin, for valuable discussions. D.R. was supported by NIH grants GM100233 and HG006399 and by NSF grant 1032255. S.S. was supported by NIH grants R01GM078598 and R01MH101244. R.D was supported by a Banting fellowship from the Canadian Institutes of Health Research.

